# Local brainstem circuits contribute to reticulospinal output in the mouse

**DOI:** 10.1101/2021.01.11.426243

**Authors:** Jeremy W. Chopek, Ying Zhang, Robert M Brownstone

## Abstract

Glutamatergic reticulospinal neurons in the gigantocellular reticular nucleus (GRN) of the medullary reticular formation can function as command neurons, transmitting motor commands to spinal cord circuits. Recent advances in our understanding of this neuron-dense region have been facilitated by the discovery of expression of the transcriptional regulator, Chx10, in excitatory reticulospinal neurons. Here, we address the capacity of local circuitry in the GRN to contribute to reticulospinal output. We define two sub-populations of Chx10-expressing neurons in this region, based on distinct electrophysiological properties and somata size (small and large), and show that these correspond to local interneurons and reticulospinal neurons, respectively. Using focal release of caged-glutamate combined with patch clamp recordings, we demonstrated that Chx10 neurons form microcircuits in which the Chx10 interneurons project to and facilitate the firing of Chx10 reticulospinal neurons. We discuss the implications of these microcircuits in terms of movement selection.

**SIGNIFICANCE STATEMENT:** Reticulospinal neurons in the medullary reticular formation play a key role in movement. The transcriptional regulator Chx10 defines a population of glutamatergic neurons in this region, a proportion of which have been shown to be involved in stopping, steering, and modulating locomotion. While it has been shown that these neurons integrate descending inputs, we asked whether local processing also ultimately contributes to reticulospinal outputs. Here, we define Chx10-expressing medullary reticular formation interneurons and reticulospinal neurons, and demonstrate how the former modulate the output of the latter. The results shed light on the internal organization and microcircuit formation of reticular formation neurons.

## INTRODUCTION

Movement is complicated. When we reach and grasp our coffee cup, we instruct our spinal cords to activate particular muscle groups in a precise temporal sequence. At the same time, we instruct it to not supinate our forearm. This combination of instructions to move and to not move are thus required for behavioural output. While the basal ganglia are involved in making these selections (Tecuapetla et al., 2016; Lee et al., 2020), it is the brainstem reticular formation that must ultimately decode these supramedullary inputs to form precise commands that descend, via reticulospinal pathways, to spinal motor circuits.

It is well established that the gigantocellular reticular nucleus (GRN) contains reticulospinal neurons that mediate posture and locomotion in the cat (Takakusaki et al., 2003; Matsuyama et al., 2004; Schepens and Drew, 2004). Neurons in supramedullary locomotor regions such as the mesencephalic (MLR) and cerebellar locomotor regions project to the GRN, where they are integrated to initiate locomotion via reticulospinal pathways that in turn activate spinal locomotor circuits (Orlovsky, 1970; Noga et al., 2003). Electrical (Jordan et al., 2008) or chemical (Noga et al., 1988) activation of glutamatergic RSNs in the GRN is sufficient for inducing locomotor activity, and inactivation (Noga et al., 2003) of this region eliminates MLR-evoked locomotion. But glutamatergic RSNs are involved in other movement as well: for example, these neurons also receive input from postural centres to ensure appropriate extensor tone during motor activities (see review:Takakusaki, 2017).

The GRN is comprised of various types of neurons, including inhibitory as well as excitatory reticulospinal neurons (Du Beau et al., 2012; Mitchell et al., 2016; Valencia Garcia et al., 2018), and is not seemingly structurally organised in a way that would facilitate the study of different neuronal types. Mouse genetic techniques have therefore been used to identify and activate (or inactivate) neurons in this region to study their roles. In recent years, neurons in the GRN that express the transcriptional regulator Chx10 have been studied (Crone et al., 2012; Bretzner and Brownstone, 2013). As in the spinal cord, these neurons are exclusively glutamatergic (Bretzner and Brownstone, 2013; Bouvier et al., 2015). Promoters for either the vesicular glutamate transporter 2 (vGluT2), or Chx10 have been used to drive cre recombinase expression for opto- or chemo-genetic experiments. Using these techniques, it has become clear that excitatory GRN neurons can modulate locomotor activity (Lemieux and Bretzner, 2019), that bilateral activation of Chx10 RF neurons can stop locomotion (Bouvier et al., 2015), and that their unilateral stimulation can lead to turning (Cregg et al., 2020) or neck movements (Capelli et al., 2017) or both (Usseglio et al., 2020). But glutamatergic neurons in this region are also involved in sleep atonia (Saper et al., 2010), suggesting that results from photoactivation experiments could be skewed by the subpopulation(s) of neurons activated. But it is likely that, similar to the heterogeneity of spinal Chx10 interneurons (e.g. Dougherty et al., 2013; Hayashi et al., 2018), Chx10 reticular formation neurons are also not homogeneous. Furthermore, opto- or chemo-genetic studies would not capture the intrinsic organisation of the GRN, and whether local circuits regulate the output – that is, whether this region is simply an integrator of inputs, or whether it has the circuitry to refine descending output.

In a step towards understanding this black box region of the brain and how it might process input from above to produce descending commands for spinal cord circuits to process, we asked whether Chx10 neurons in the GRN form local microcircuits. We identified two clear subsets of these neurons, only one of which (comprised of neurons with large somata) is reticulospinal. Then, using focal release of caged glutamate, we demonstrated the internal architecture whereby smaller Chx10 neurons project to and facilitate the firing of the larger Chx10 neurons. We propose that these microcircuits tune descending commands and thus facilitate movement.

## MATERIALS AND METHODS

### Animals

All animal procedures were done at Dalhousie University, approved by the University Committee on Laboratory Animals and conform to the standards of the Canadian Council for Animal Care. Both male and female Chx10::eGFP were used in all experiments.

### Electrophysiology

#### Slice preparations

Whole-cell patch clamp recordings were made from visualized Chx10 neurons from the gigantocellular reticular nucleus (GRN) of the rostral medulla of P9-P14 Chx10e:GFP mice (Bretzner and Brownstone, 2013). Mice were anesthetized by intraperitoneal injection of a ketamine (60 mg/kg) and xylazine (12 mg/kg). After loss of their righting reflex, mice were cooled on ice and decapitated. The brainstem was immediately removed and secured to an agar block with a small amount of adhesive and transferred to ice-cold oxygenated solution (3.5mM KCL, 35mM NaHC03, 1.2mM KH2P04, 1.3mM MgS04, 1.2mM CaCl2, 10mM glucose, 212.5mM sucrose, 2mM MgCl2, ph7.4) for sectioning. Two, 300-350 um sections of the medulla containing the rostral GRN were sectioned on a vibratome (Leica VT1200S, Leica) per mouse and transferred to a warm oxygenated (95% oxygen, 5% carbon) aCSF solution (111 mM NaCl, 3.08 mM KCl, 11 mM Glucose, 25 mM NaHC03, 1.25 mM MgS04, 2.52 mM CaCl2, 1.18 mM KH2p04, pH 7.4) to recover for a minimum of 30 minutes.

#### Whole-cell patch-clamp recordings and stimulation

Slices were transferred to a recording chamber mounted on a Zeiss AxioExaminer microscope and perfused with oxygenated room-temperature aCSF. Cells were visualized using a 20x wide aperture (1.2nA) water-immersion objective lens, a CCD camera (CoolSNAP EZ CCD Camera, Photometrics, Arizona, USA) and Slidebook 6.0 software (Intelligent Imaging Innovations, Colorado, USA, RRID:SCR_014300).

Whole-cell patch-clamp recordings were acquired in current-clamp (IC) mode using a Multiclamp 700B amplifier (Molecular Devices, California, USA, RRID:SCR_014300). Recordings were loss pass filtered at 10 kHz (IC), and acquired at 25 kHz with CED Power1401 AD board and Signal software (CED, Cambridge UK, http://ced.co.uk/us/products/sigovin).

Recording pipettes were filled with a solution containing in mM: K-gluconate, 128; NaCl, 4; CaCl2, 0.0001; Hepes, 10mM, glucose, 1mM; Mg-ATP, 5; and GTP-Li, 0.3, pH 7.2, lucifer yellow dilithium salt (0.4 mg/ml, Thermo Fisher Scientific, Cat# A-5750, RRID:AB_2314410), and neurobiotin (1mg/ml, Vector Laboratories, Cat# SP-1120-20, RRID: AB_2536191), and had resistances of 4-6 MΩ. Lucifer yellow allowed for the immediate visualization of the soma and dendrites of the patched cell. This allowed us to avoid photostimulating processes of the recorded cell whilst stimulating nearby presynaptic cell bodies. Neurobiotin allowed for *post-hoc* confirmation of the identity of the recorded cell and that axons or dendrites of the post-synaptic cell were not stimulated when photostimulating pre-synaptic cell bodies by comparing images of the region of interest (ROIs) stimulated (see below) to the confocal image of the biotin-filled post-synaptic cell (n=10).

In the initial set of experiments, basic and rhythmic firing properties of visually identified “large” and “small” Chx10 neurons in the medulla GRN were subjected to various depolarizing and hyperpolarzing current pulses. All properties were collected while holding the cell at −60 mV. Input resistance was collected as the average response of the cell to repetitive (minimum 20 sweeps), small hyperpolarizing pulses (−10pa, 100ms). Rheobase, defined as the minimum current to elicit an action potential 50% of time was collected from incremental 1 pA depolarizing current steps. Voltage threshold defined as the membrane potential at which depolarization increased at ≥10 V/s was determined from the first spiking response during rheobase. Frequency-current (F-I) plots were obtained by applying 1 second depolarizing current pulses with incremental increase of 10-20 pA. The steady state firing frequency was determined by counting the number of spikes during the 1 second pulse. The instantaneous firing frequency was collected by taking the average interspike interval (1/average interspike interval) of the first 3 spikes for each current step. Sag potential and post inhibitory rebound firing was collected during 1 second hyperpolarizing current pulses with increases of −10 to −20 pA. Sag potential was collected as the difference in the membrane potential at maximum minus steady state change during the hyperpolarizing pulse. All sag calculations were conducted at a steady state hyperpolarization of −100 mV.

### Holographic photolysis of caged glutamate

Following the initial set of experiments to characterize the electrophysiological properties of large and small Chx10 neurons, we sought to determine the connectivity patterns of Chx10 neurons using a photo-stimulation protocol as we have previously described. These experiments only proceeded when membrane potential was stable (i.e. did not fluctuate more than 5 mV during a 5-minute period). MNI-caged-L-glutamate (2.5mM, Tocris, Ontario, Canada, Cat# 1490) was perfused in warm aCSF at a rate of 2ml/min. Holographic photolysis of MNI-glutamate was performed using a 405 nm laser directed through a Phasor spatial light modulator (SLM) system (Intelligent Imaging Innovations, Colorado, USA), stimulating regions of interest (ROI) as controlled by Slidebook software. Prior to photostimulating ROIs, direct photostimulation of the patched Chx10 neuron was performed to determine the intensity and duration of the pulse which elicited a single action potential. Ideal intensity and pulse duration for small Chx10 neurons was 3.5mW and 800-1000 μs and for large Chx10 neurons was 7mW and 1500 μs. If longer durations were necessary, the slice was not used as it was deemed to be unhealthy. ROIs were selected as identified neuronal cell bodies under fluorescence, with the ROI blanketing the soma. Once ROIs in a single focal plane were selected, sequential photostimulation of the ROIs was carried out while recording from the post-synaptic neuron. Post-synaptic cells were held between −60mv and −50mv for the duration of the experiment (which could exceed one hour). Photomicrographs for each ROI were taken to map the location of each cell relative to the post-synaptic cell recorded. In stable preparations, once a connection was established, a second electrode was used to record from the connected pre-synaptic cell to determine if the connection was bi-directional. As the connected cells were often in close proximity (100-200 μm), the gigaseal of the patch-clamp was often lost on one of the two cells, but loose patch recordings were obtained, confirmed by recording responses from direct photostimulation of the cell body.

### CTB injections

The fluorescent retrograde tracer cholera toxin subunit B (CTB) was used to label reticulospinal neurons. Briefly, using aseptic surgery with 3-4% isoflurane, an incision over the cervical or lumbar spine was made, the lamina over C6 or L2 was removed, and a small incision in the dura was made to expose the spinal cord. Mice were placed in a stereotaxic frame to stabilize the vertebral column, and, using a micromanipulator a glass pipette containing 1 μL of CTB was advanced to the surface of the spinal cord, following which the tip was slowly lowered 500 μm into the L2 spinal cord. CTB was injected over a 10 minute period and the pipette remained in place for 5 minutes after injection, after which, the electrode was removed and a similar injection with a new glass electrode was made on the contralateral side of the spinal cord. The mice were then sutured and monitored while they recovered. No deficits or motor impairments were seen as mice recovered. At day 14 after injection, the mice were anesthetized and transcardially perfused with phosphate buffer solutions followed by 4% paraformaldehyde in 0.1M phosphate solution (PBS). Tissue was harvested, post-fixed in 4% paraformaldehyde overnight, and subsequently cryoprotected in 30% sucrose. Tissue was sectioned using a cryostat and mounted on glass slides for immunohistochemistry.

### Immunohistochemistry and imaging

Upon completion of the electrophysiological recordings, slices were incubated in 4% paraformaldehyde for 1 hour at room temperature followed by three, 15 minute washes in 0.1% PBS-T. Slices were incubated at 4°C overnight in goat anti-GFP primary antibody (1:2500, Abcam Cat# AB6673, RRID: AB_305643) followed by a four-hour incubation period in donkey anti-goat Alexa Fluor 488 (1:500, Abcam Cat# AB150105, RRID: AB_2732856) and Alexa Fluor 647-conjugated streptavidin (1:500, ThermoFisher Scientific, Cat# S-21374. RRID: AB_2336066). Images were obtained using a Zeiss LSM 510 upright confocal microscope. Images were compared to photos obtained during the electrophysiology recordings in Slidebook to confirm that the ROIs did not overlap with visually identifiable dendrites or axons.

For CTB-labeled tissue, sections were incubated with primary goat anti-GFP (1:500, Abcam, Cat# AB5449, RRID: AB_304896) and sheep anti-Neun (1:1000, Abcam Cat# AB177487. RRID: AB_2532109) antibodies diluted in PBS containing 0.1% Triton X-100 (PBS-T) for 24 hours at 4 °C. Sections were then washed three time for 10 mins each in 0.1% PBS-T followed by incubation with appropriate secondary antibodies conjugated to Alexa 488, Alexa 555 and Alexa 637 (1:500 Molecular Probes) for 3 hours. Sections were then washed three times for 10 mins each in PBS, mounted in vectashield (Vector Laboratories) and coverslipped. Epifluorescent images were acquired with a Zeiss Axioplan inverted microscope. Images were processed using ImageJ software (NIH, USA). Cell areas (μm^2^) were measured at optical planes that included the nucleus, and were calculated using surface area function in ImageJ

### Experimental design and statistical analysis

Data are presented as mean ± SD. As is common for discovery experiments, no statistical method was used to predetermine sample size, and no randomization or blinding procedures were used. Statistical analyses were done using Sigma-Plot (v14.0, Systat software, California, USA). Un-paired Student’s t test were used for all comparisons except for frequency of observed sag potentials, in which Fisher Exact test was used. Statistical significance was set at p < 0.05.

## RESULTS

### Chx10 neurons have distinct electrophysiological properties correlated to their size

As intrinsic neuronal electrophysiological properties play a critical role in producing circuit activity (Getting, 1989), we first sought to characterize these properties of medRF Chx10 neurons, focusing on those in the GRN. We performed whole-cell patch-clamp recording from Chx10 neurons in P9-P14 mice to determine if there are subpopulations of Chx10 neurons with distinct electrophysiological properties. All neurons were held at −60 mV to have a uniform baseline when measuring their responses to injected current.

We first characterized the spike patterns of Chx10 neurons in response to sustained suprathreshold current steps. Of the 60 neurons we recorded, 48 demonstrated sustained firing in response to current injection, whereas the remaining 12 fired only a single spike and were not analysed further. Of the 48 neurons with sustained firing, two distinct patterns of firing emerged. The first pattern (n=26) consisted of trains of action potential with minimal spike frequency adaptation (SFA) and lower initial gain in the F/I relationship (Figures 1A &1B). The second pattern (n=22) consisted of high initial firing rate followed with pronounced spike frequency adaptation. The average gain for the initial firing (first 3 spikes) was 19 ± 18 Hz / 10 pA for the first pattern of firing Chx10 neurons and 36 ± 28 Hz / 10pA (P =0.02) for the second type of firing Chx10 neurons (Figure 1B). There was no significant difference in the average gain for the steady firing (last 3 spikes), with average gains of 9 ± 9 Hz / 10pA s and 12 ± 9 Hz / 10 pA respectively (Figure 1B, P = 0.08). Furthermore, the second type of firing Chx10 neurons demonstrated significantly more SFA compared to the first type of Chx10 neurons (43 ± 12 % vs 26 ± 20 % decrease in firing rates, respectively, P =0.028). Based on these distinct firing patterns, we termed these neurons Type 1 and Type 2 Chx10 neurons, and then analysed their corresponding electrophysiological properties (Figure 1C; see below).

**Figure 1.**
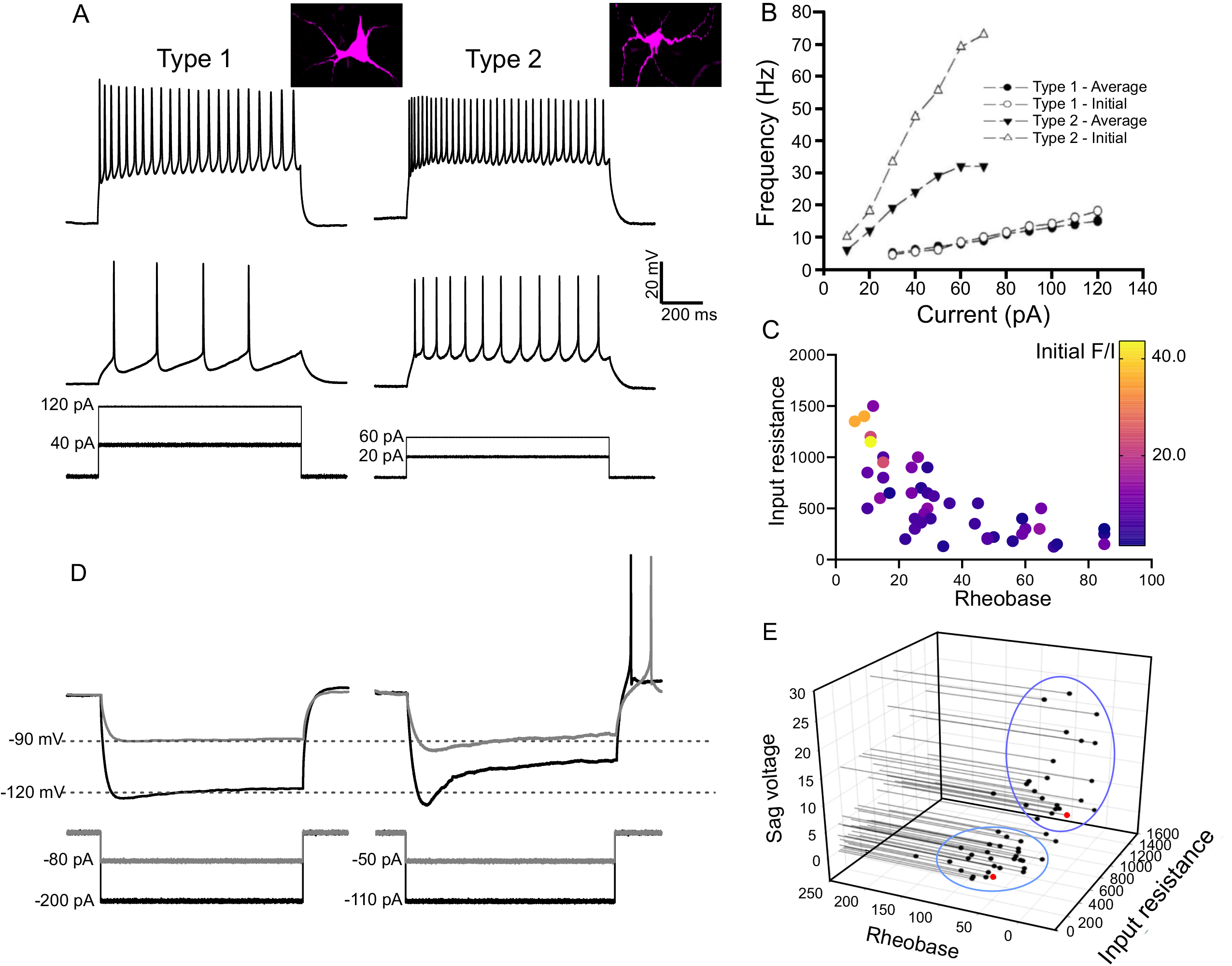
Chx10 neurons have two distinct types of electrophysiological properties. **A)** In response to incremental depolarizing current steps, Type 1 Chx10 neurons demonstrate minimal gain in their firing rate with little spike frequency adaptation, whereas Type 2 Chx10 neurons demonstrate a high initial firing rate with SFA. **B)** Frequency-current relationship plots of the initial and average firing rates for Type 1 and Type 2 Chx10 neurons. **C)** Relationship of initial f-I slope with input resistance and rheobase. Colour reflects the f-I gain (Hz/10 pA; see sidebar), as does size. **D)** In response to incremental hyperpolarizing current steps, Type 1 Chx10 neurons demonstrate minimal sag and no PIR, whereas Type 2 Chx10 neurons demonstrate substantial Sag and PIR following termination of the hyperpolarizing pulse. **E)** 3D plot showing the sag voltage, rheobase and input resistance recorded from each Chx10 neuron. Blue circles showing separation of Type 1 and Type 2 Chx10 neurons based on their plotted electrophysiological properties. The neurons indicated by the 2 red dots are shown in Figure 5.

The two types of Chx10 neurons also demonstrated differences in their responses to hyperpolarizing current injection to steady state voltages of −100 mV. Of the 26 Type 1 Chx10 neurons, 10 had sag potentials, in response to a hyperpolarizating step to −100 mV, and these were of small amplitude (3.8 ± 5.1 mV). None of these neurons responded with post-inhibitory rebound (PIR) upon termination of this step (Figure 1D). Conversely, all Type 2 Chx10 neurons (n=22) had a large sag potential (21.4 ± 7.5 mV, P < 0.001) in response to hyperpolarization (Fisher Exact P < 0.001), and PIR responses consisted of a single or several action potentials at pulse termination (Figure 1D). To summarize, we found that Type 1 Chx10 neurons demonstrated: 1) lower gains in their initial firing F/I relationships, 2) minimal SFA, 3) no sag potentials, and 4) no PIR, whereas Type 2 Chx10 neurons demonstrated: 1) higher gains in their initial firing F/I relationships, 2) significant SFA, 3) sag potentials, and 3) PIR responses. Based on these two distinct types of firing patterns in response to depolarizing and hyperpolarizing current injections, we proceeded to compare passive electrical properties of these two types of Chx10 neurons.

Both types demonstrated similar resting membrane potentials (−46.8 ± 5.4 mV and −46.2 ± 6.6 mV, Type 1 and 2 respectively, P = 0.73) and spike thresholds (−45.2 ± 6.4 mV and −43.4 ± 3.7 mV, Type 1 and 2 respectively, P = 0.23). However, there was a significant difference in input resistance with Type 1 demonstrating an average input resistance of 380 ± 245 MΩ while Type 2 had an average input resistance of 980 ± 480 MΩ (P < 0.001). Correspondingly, Type 1 Chx10 neurons were less excitable with an average rheobase of 56.0 ± 40.8 pA, compared to Type 2 Chx10 neurons which had an average rheobase of 21.0 ± 7.5 pA (P < 0.001). These 3 properties – input resistance, rheobase, and sag – together led to a separation between Type 1 and Type 2 Chx10 neurons (Figure 1E).

During these recordings, it became visually evident that Type 1 Chx10 neurons were larger than Type 2 Chx10 neurons (Figure 1A inset). This observation matched their different rheobases and input resistances. Furthermore, we injected biotin into patched neurons and analyzed their soma sizes and the number of initial dendritic branches (n=5 for each type). It was clear that Type 1 Chx10 neurons are significantly larger (385 ± 48 μm vs 148 ± 45 μm, P <0.001) and have more primary dendrites (3.4 ± 0.5 vs 1.4 ± 0.5, P = 0.004) than Type 2 Chx10 neurons. The difference in soma size fits with a bimodal distribution of Chx10 soma sizes (vide infra). We therefore refer to these two types of GRN neurons as Large Chx10 neurons and Small Chx10 neurons, and next sought to determine their connectivity patterns.

### Large Chx10 neurons are reticulospinal

As Chx10 neurons have previously been shown to be reticulospinal (Bretzner and Brownstone, 2013 & Bouvier et al., 2015) we sought to determine whether both subpopulations project to the spinal cord. We initially targeted the L2 spinal cord, where we injected CTB bilaterally for retrograde labeling in adult Chx10:eGFP mice (n=3, Figure 2A). After a two week recovery period, we found, on average, 63 (range 35-82) CTB+ GFP^ON^ neurons/mouse in the GRN (Figure 2A). This represented 8-11% of all Chx10^ON^ neurons in the GRN, similar to that reported in neonatal mice (Bretzner & Brownstone, 2013).

**Figure 2.**
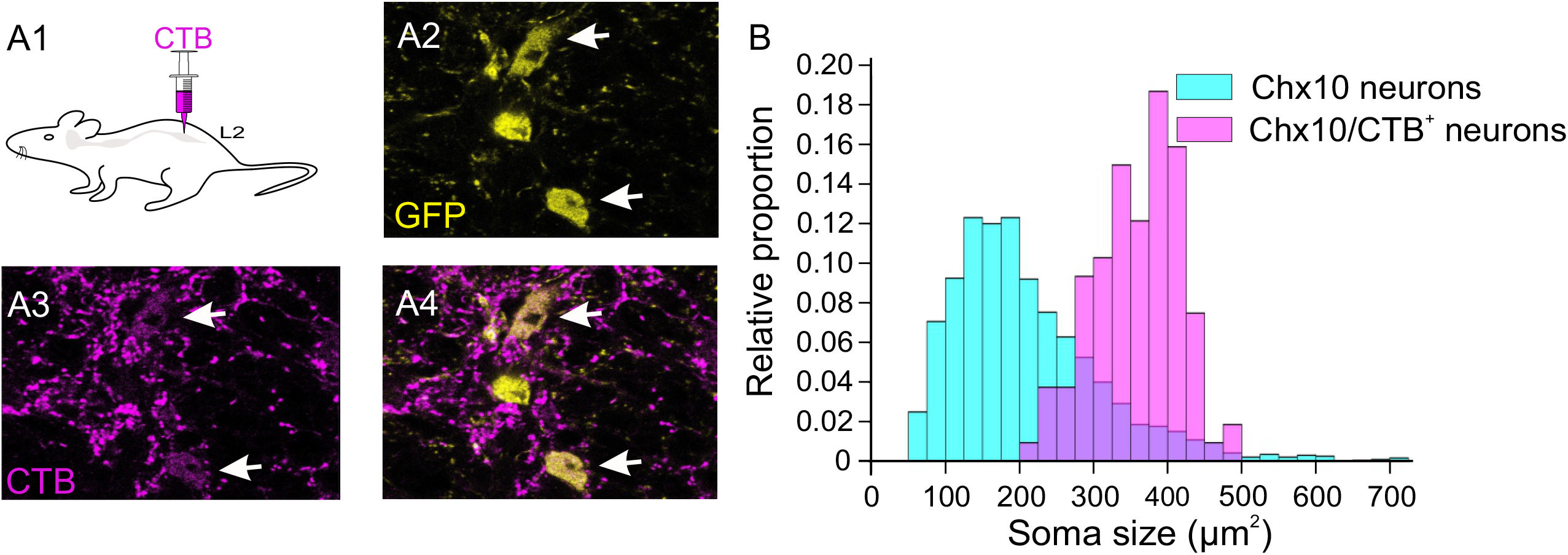
Large Chx10 neurons are reticulospinal neurons. **A1)** Schematic of experimental procedure in which CTB is injected bilaterally in the second lumbar segment to label reticulospinal neurons. **A2)** Image of GFP positive Chx10 neurons in the gigantocellular region of the medulla. **A3)** Image of CTB labelling of terminals and cell bodies in the same section as A2. **A4)** Overlap of GFP and CTB, with two Chx10 neurons positive for CTB labelling (arrows), and one smaller one that is not. **B)** Overlaid cumulative plots demonstrating the relative distribution of all labelled GFP Chx10 neurons in the GRN (light blue), as well as the relative distribution of CTB positive Chx10 neurons (pink). Note that although the frequency of large Chx10 cells is lower, it is these large cells that are reticulospinal.

As a population, Chx10 soma sizes ranged from 80 – 610 μm^2^ (n= 1925, from 3 mice mean: 211 ± 105 μm^2^). In contrast, the range of average soma sizes of CTB positive Chx10 neurons was 240 – 540 μm^2^ (mean: 360 ± 60 μm^2^; p <0 .01)), corresponding with larger Chx10 neuronal soma sizes seen in the frequency distribution (Figure 2B), indicating that the Large Chx10 neurons are reticulospinal. We found no evidence that the Small Chx10 neurons project to the lumbar spinal cord.

To investigate whether Small Chx10 neurons project to the rostral spinal cord, we injected CTB in the C7-C8 cervical segments (n=3) and quantified the number of CTB+ Chx10^ON^ neurons in the medRF GRN as above. We found a similar distribution of CTB+ Large Chx10 neurons and no CTB+ Small Chx10 neurons. Thus, Large Chx10 neurons are reticulospinal whereas Small Chx10 neurons are not. We will therefore refer to the Large Chx10 neurons as Chx10 reticulospinal neurons, and the small ones as Chx10 interneurons.

## CONNECTIVITY PATTERNS OF CHX10 NEURONS

To determine whether the two populations are synaptically connected with each other, we used brain stem slice preparations to record from 30 Chx10 neurons (15 of each type), while photo-activating caged glutamate on over 120 others, establishing 60 connections. We targeted Chx10 neurons based on their size to ensure that we recorded a sufficient number of each type. Once entering whole cell mode, we first characterized the neurons based on their electrophysiological properties (see above), and found a 96% (29/30) concordance rate between their properties and the predicted type based on size. That is, if we treat our initial data from 48 neurons as a “training” set, then in this “test” set of 30 neurons, we picked their electrophysiological properties correctly in 29. Therefore, we were confident that we could visually classify neurons as interneurons or reticulospinal neurons for photostimulation. As can be appreciated by the distribution of cell sizes (Fig 2B), there were many more Chx10 interneurons than reticulospinal neurons. We found that Chx10 interneurons form synapses with both Chx10 interneurons and Chx10 reticulospinal neurons. Conversely, we did not find evidence that Chx10 reticulospinal neurons form synapses with other Chx10 reticulospinal neurons, and connections to Chx10 interneurons were infrequent, although in a subset of pairs, we found bi-directional connectivity between Chx10 interneurons and Chx10 reticulospinal neurons. We describe these findings below.

### Chx10 interneurons form synapses with Chx10 interneurons

After ensuring that photostimulation produced action potentials in Chx10 interneurons by directly stimulating the recorded cell (Fig 3A), we asked whether Chx10 interneurons connect to each other. In 15 Chx10 interneurons (average soma size 176 ± 40 um), 20 connections were found by photostimulation of 70 Chx10 interneurons (Figure 3B). In all 20 connections, the post-synaptic responses were subthreshold, with differing sizes of excitatory post synaptic potentials (EPSPs) arising from different presynaptic neurons (Figure 3C). In some cases, temporal summation with repetitive stimulation was evident (Fig 3C). The number of connected presynaptic neurons ranged from 1 to 3 (mean 1.7 ± 0.9), with EPSP amplitudes ranging from 4.0-12.5 mV (mean 8 ± 3 mV).

**Figure 3.**
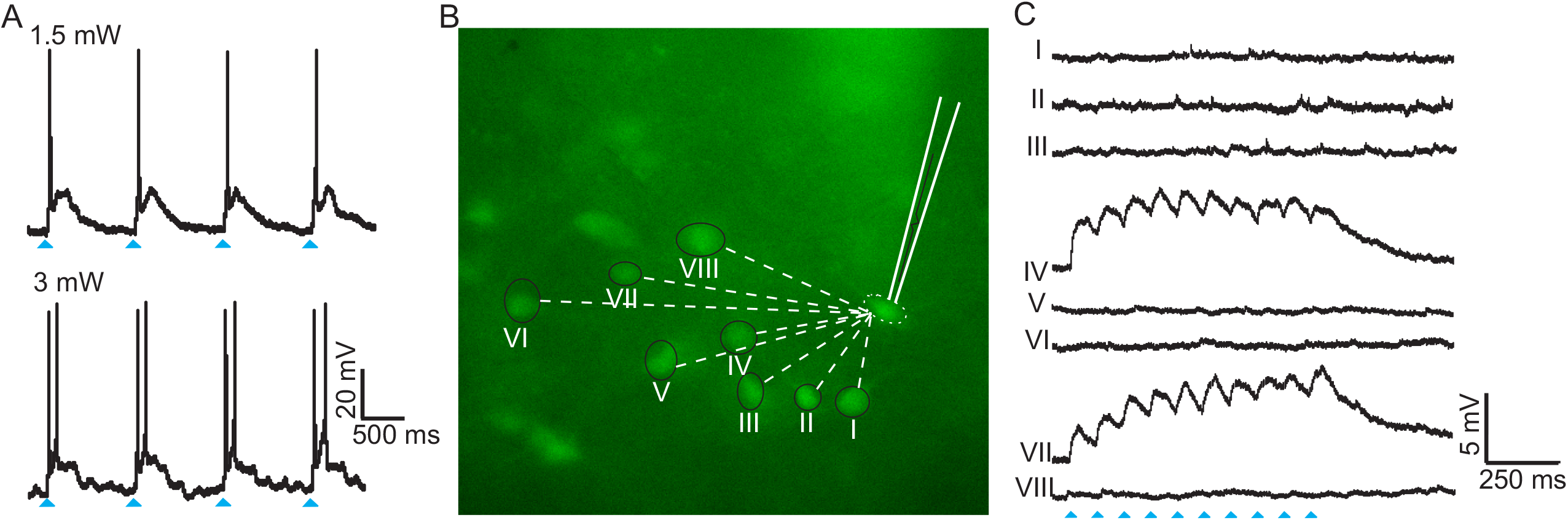
Chx10 interneurons form synapses with other Chx10 interneurons. **A)** Direct holographic stimulation over patched cell results in single or multiple action potentials, dependent on the laser output. **B)** Fluorescence image of Chx10::eGFP positive neurons in the rostral GRN. Synaptic connectivity was determined by stimulating visually identified Chx10 interneurons and recording post-synaptic potentials in the patched Chx10 interneuron. **C)** Post synaptic responses from stimulation of Chx10 interneurons with corresponding numbers in panel B. In this example, stimulation (trains) of neurons IV and VII elicited EPSPs, whereas stimulation of the other Chx10 neurons did not. Blue arrows indicates laser on. Example from n= 20 connections.

### No apparent connectivity between Cxh10 reticulospinal neurons

We next turned to Chx10 reticulospinal neurons, and asked whether they form synapses with each other. Due to the large size and high rheobase of Chx10 reticulospinal neurons, higher photostimulation intensities were needed to generate action potentials in response to the majority of the individual stimuli (Figure 4A). During recordings in 7 Chx10 reticulospinal neurons, we photostimulated a total of 40 other Chx10 reticulospinal neurons but were unable to establish connectivity: there were no responses.

**Figure 4.**
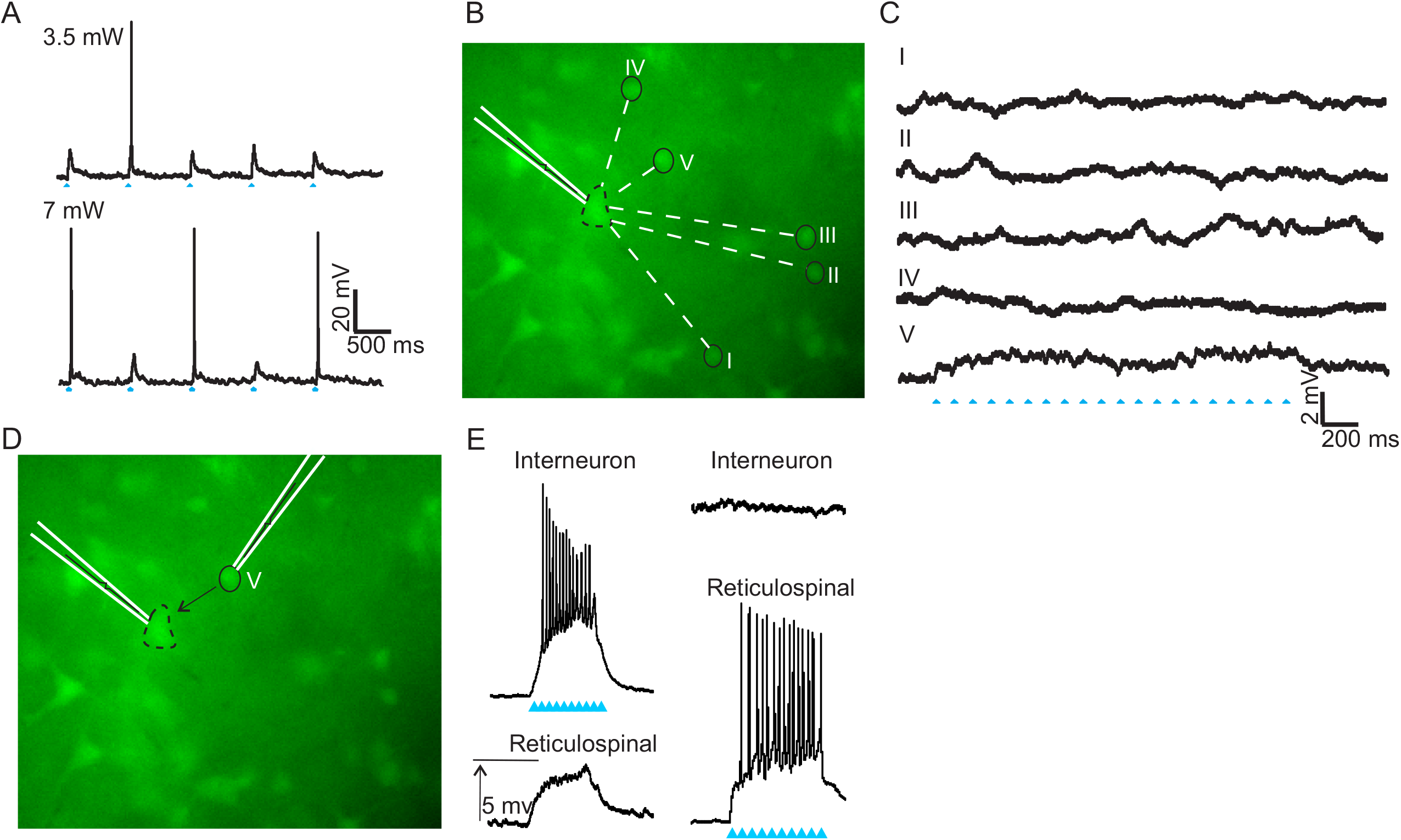
Chx10 interneurons form synapses with Chx10 reticulospinal neurons. **A)** Direct holographic stimulation of patched RSN results in EPSPs or single action potentials dependent on laser output. **B)** Fluorescence image of Chx10::eGFP positive neurons in the rostral medulla. Synaptic connectivity was determined by stimulating visually identified Chx10 interneurons and recording post-synaptic potentials in the patched Chx10 RSN. **C)** Post synaptic responses from stimulation of numbered Chx10 interneurons. In this example, spot V elicited EPSPs from a train of laser stimuli, whereas stimulation of the remaining Chx10 neurons did not elicit a response. **D)** After establishing a synaptic connection, a second electrode was used to patch and recorded responses in the Chx10 interneuron (spot V). **E)** Photostimulation of the Chx10 interneuron resulted in multiple actional potentials and post synaptic EPSP in the Chx10 reticulospinal neuron. **F)** Photostimulation of the Chx10 reticulospinal neuron elicited multiple action potentials but did not elicit a response in the Chx10 interneuron. Blue arrows indicate laser on. Example from n= 32 connections.

### Chx10 interneurons form synapses with Chx10 reticulospinal neurons

We next turned to investigating connectivity between the two types of neurons. To study connectivity from Chx10 interneurons to Chx10 reticulospinal neurons, we photostimulated 75 Chx10 interneurons whilst recording Chx10 reticulospinal neurons (n=15, soma size 421 ± 84 um), and found a total of 32 connections in 14 neurons (Figure 4B). In all 32 connections, the post-synaptic responses were subthreshold, resulting in small post synaptic EPSPs (3.2 ± 1.2 mV, Figure 4C, presynaptic neuron V). The number of connections ranged from 1 to 6 presynaptic neurons, with an average of 2.5 ± 1.3 connections per patched Chx10 reticulospinal neuron. Conversely, in 5 Chx10 interneurons, we photostimulated a total of 25 Chx10 reticulospinal neurons but were unable to establish any connections, suggesting that connectivity between the two populations is unidirectional.

In a subset of experiments that showed connectivity from a Chx10 interneuron to a Chx10 reticulospinal neuron (n=7), a second patch electrode was used to successfully record the pre-synaptic Chx10 interneuron, although this often resulted in the loss of the initial giga seal on the postsynaptic Chx10 reticulospinal neuron. (Figure 4D). In these experiments, action potentials generated in the Chx10 interneurons by direct photostimulation led to EPSPs in the Chx10 reticulospinal neurons as expected (Figure 4E). In 5/7 of these experiments, photostimulation of Chx10 reticulospinal neurons that resulted in generating action potentials in the Chx10 reticulospinal neurons did not elicit post-synaptic responses in the Chx10 interneurons, supporting that these connections are uni-directional.

In two paired recording experiments, however, bi-directional connectivity was found using current injections into the identified presynaptic Chx10 interneuron and Chx10 reticulospinal neurons (Figure 5A, B), Incremental depolarizing current steps injected in Chx10 interneurons (Figure 5C) that generated action potentials led to EPSPs and action potentials in Chx10 reticulospinal neurons (light grey and black, figure 5C) at low (light red, figure 5C) and high (dark red, figure 5C) current injections, respectively. Similarly, current injection in Chx10 reticulospinal neurons (figure 5D) that generated action potentials also generated EPSPs and an action potential (light and dark red, figure 5D) in the Chx10 interneuron at low (light grey, figure 5D) and high current injections (black, figure 5D), respectively (Figure, demonstrating that in at least some cases, connectivity is bi-directional.

**Figure 5.**
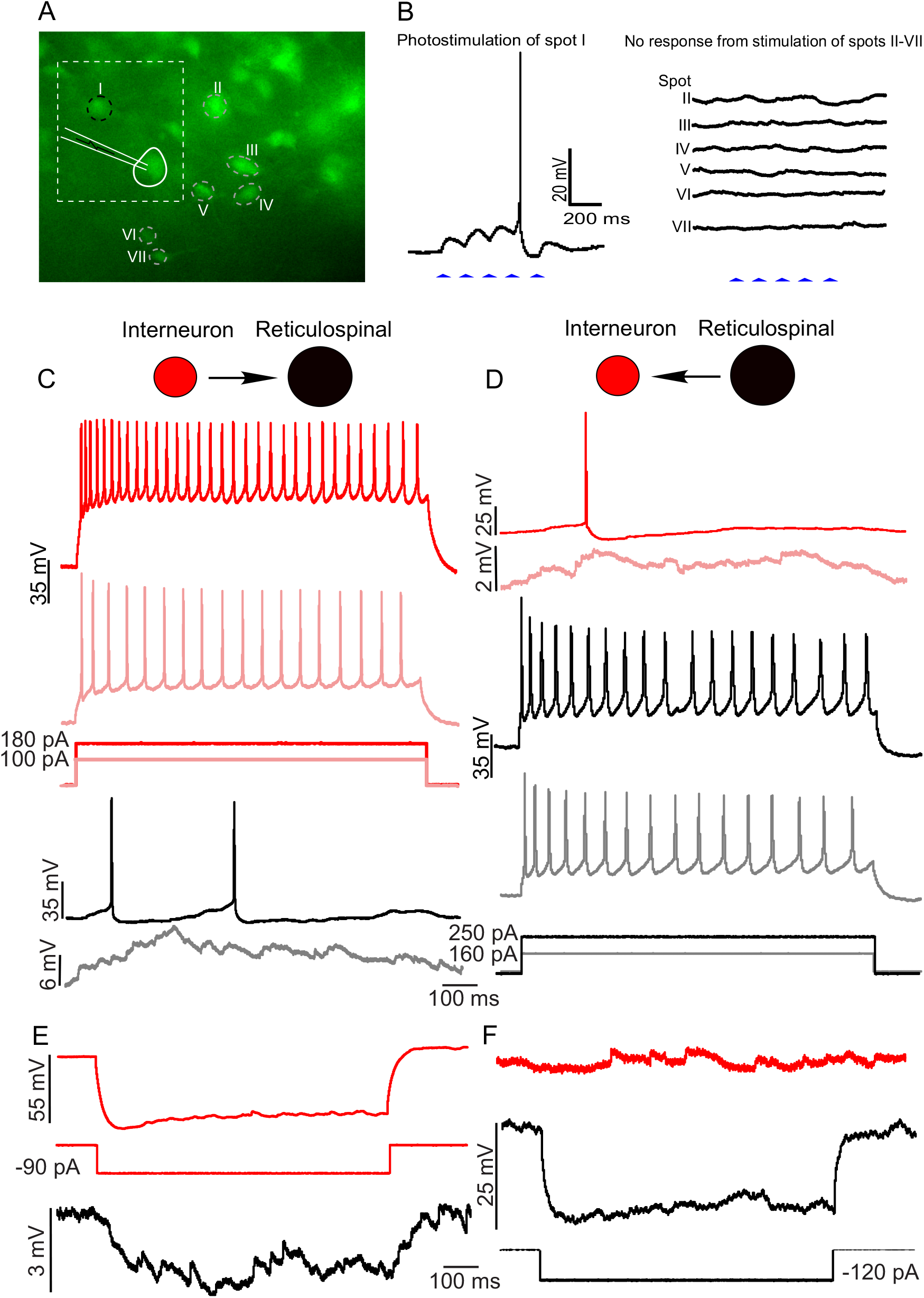
Bi-directional connectivity between a sub-set of Chx10 interneurons and Chx10 reticulospinal neurons. **A)** Fluorescence image of Chx10::eGFP positive neurons in the rostral GRN. Synaptic connectivity was determined by stimulating visually identified Chx10 interneurons whilst recording post-synaptic potentials in the patched Chx10 RSN. **B)** Photostimulation of neuron 1 elicited EPSPs that summated to produce an action potential in the RSN. After establishing this synaptic connection, a second electrode was used to patch and record responses in the Chx10 interneuron (neuron 1). Note that electrophysiological properties of these 2 neurons are shown by the red dots in Figure 1E. **C)** Depolarizing current injections in the Chx10 interneuron resulted in action potentials in the Chx10 interneuron (red) and EPSPs and action potentials in the Chx10 RSN (black) at low and high current intensities, respectively. **D)** Similarly, depolarizing current injections in the Chx10 RSN resulted in action potentials in the Chx10 RSN, and EPSPs and action potentials in the Chx10 interneuron at low and high current intensities, respectively. **E)** Hyperpolarizing current injections in the Chx10 interneuron resulted in a small hyperpolarizing response in the Chx10 RSN. **F)** Hyperpolarizing current injections in the Chx10 RSN did not elicit a detectable response in the Chx10 interneuron. Note that the RSN could only be stably hyperpolarized by ~25 mV. Example from n=3 dual patch recordings.

Given this bidirectionality, and given that Cx36 is widely expressed in the medRF GRN (Westberg et al., 2009; Martin et al., 2011), we sought to determine if these connections could have an electrical component, by delivering hyperpolarizing current injections. When Chx10 interneurons were hyperpolarized, there was indeed a small hyperpolarizing response in Chx10 reticulospinal neurons (Figure 5E), suggesting an electrical component to this connection. However, hyperpolarizing current injections in Chx10 reticulospinal neurons failed to elicit a concomitant response in Chx10 interneurons (Figure 5F). The failure to evoke a hyperpolarizing response in the Chx10 interneuron could be a result of a rectifying electrically coupled connection (Rash et al., 2013 & O’Brien, 2014) or could reflect that we were unable to hyperpolarize the Chx10 reticulospinal neuron sufficiently to detect a response in the Chx10 interneuron, as Chx10 reticulospinal neurons became unstable during sustained hyperpolarization.

### Chx10 interneurons contribute to repetitive firing behaviour of Chx10 reticulospinal neurons

To determine the degree to which inputs from Chx10 interneurons impact the firing behavior of Chx10 reticulospinal neurons, we photostimulated Chx10 interneurons while inducing firing in Chx10 reticulospinal neurons (n=3). After a connection was established, Chx10 reticulospinal neurons were subjected to ramp current injections alone and during photostimulation of a connected Chx10 interneuron (Figure 6A). At its peak, photostimulation could double the firing rate of the Chx10 reticulospinal neuron, increasing peak firing rate from 10 ± 2 Hz to 22 ± 3 Hz. Similar increases in firing rate were seen when Chx10 interneurons were photostimulated during rectangular depolarizing current (Figure 6B). The average firing rate of 7 ± 2 Hz doubled to 15 ± 4 Hz during photostimulation of the connected Chx10 interneuron. Interestingly, these induced firing rates are higher than those seen with maximal current injections into the recorded neurons (Figure 1, F/I plots). The physiology underlying these photostimulation-induced firing patterns – acceleration during the stimulus and slow return to baseline – is unclear, but the patterns are similar to those seen with activation of persistent currents in neuronal dendrites (Heckman et al., 2003).

**Figure 6.**
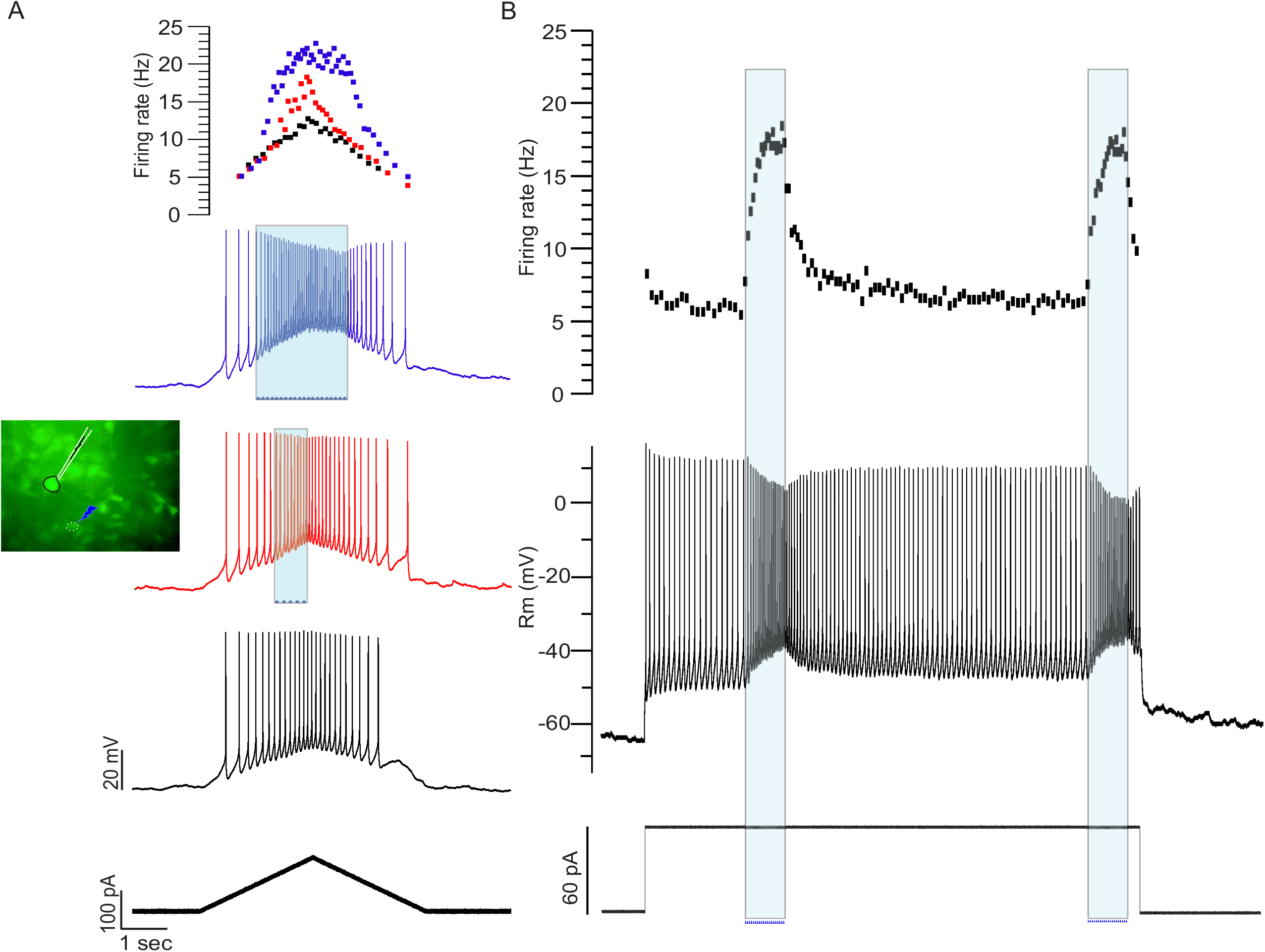
Chx10 interneurons mediate the excitability of Chx10 reticulospinal neurons. **A)** A Chx10 RSN (inset showing patched neuron) was subjected to ramp current injections (bottom trace) without (black trace) and with (red and blue traces) photostimulation of a connected Chx10 interneuron (inset, blue arrow). In the middle trace (red), the photostimulation began later and lasted a shorter period of time than in the top trace (blue). The firing rate of the RSN was plotted with and without photostimulation (top trace). B) Same connected Chx10 interneuron photostimulated (shaded blue) during sustained depolarizing current injection into the Chx10 RSN, with the resulting firing rate shown in the top trace. Example from n= 3 cells.

## DISCUSSION

Motor circuits rely on local interneuronal processing to coordinate and regulate movement. To address the organization of neurons in the medullary reticular formation, a site critical for movement, we used whole cell patch clamping to record the electrophysiological properties of Chx10^ON^ neurons. We identified two subtypes of Chx10 neurons in the GRN, one with large somata that is reticulospinal, and the other with smaller somata that functions as local excitatory interneurons. We then determined their local connectivity patterns using holographic photolysis of caged glutamate. We found that Chx10 interneurons commonly formed connections with other Chx10 interneurons as well as with Chx10 reticulospinal neurons, and that these local interneurons mediate the excitability of the reticulospinal neurons. We propose that GRN Chx10 interneurons are involved in processing and regulating the input:output functions of Chx10 reticulospinal neurons.

### Subtypes of GRN Chx10 Neurons

It is not surprising that the GRN is comprised of heterogeneous neuronal types. Electrophysiological analysis of neurons in the reticular formation of macaques has similarly demonstrated several (likely 4) different clusters of neurons based on their firing properties (Zaaimi et al., 2018). Only one of these clusters was thought to be reticulospinal, suggesting the presence of other local or ascending neurons. Taken together with our data, it seems likely that distinct electrophysiological phenotypes in the reticular formation reflect different targets and functions of the recorded neurons.

Chx10, or Vsx2, is a transcriptional regulator that is expressed in some glutamatergic neurons in both the spinal cord and brain stem, as well as in other sites (e.g. retina). In the spinal cord, it is expressed in V2a neurons – the excitatory neurons of the cardinal V2 class. It is clear that spinal V2a neurons, like other cardinal classes (e.g. V0s – Pierani et al., 2001; Zagoraiou et al., 2009; Talpalar et al., 2013; V1s – Bikoff et al 2016; V2s – Lundfald et al., 2007; Peng et al., 2007; V3s-Zhang et al., 2008; Borowska et al., 2013; Chopek et al., 2018) are not homogeneous: for example, a subset of V2a interneurons expresses Shox2 and may regulate burst variability and pattern during locomotion (Dougherty et al., 2013). And only a subset of V2a neurons in the cervical spinal cord projects to the brain stem (Hayashi et al., 2018). Likewise, we had previously demonstrated heterogeneity of Chx10^ON^ neurons in the reticular formation (Bretzner & Brownstone 2013). But we now add to this by demonstrating that there are two clear electrophysiological phenotypes of Chx10^ON^ neurons in the GRN, and these phenotypes correlated with whether or not the neurons had reticulospinal projections.

### Functional Diversity of Chx10 GRN Neurons

While we have shown that we can readily classify Chx10^ON^ GRN neurons into two populations, there are likely many more sub-classes. In the spinal cord, it is increasingly clear that genetically defined neuronal populations are not homogenous and that subtypes have different electrical properties, connectivity and function (Bikoff et al., 2016). Some GRN Chx10 neurons are involved in respiration (Crone et al., 2012; Romer et al., 2017; Jensen et al., 2019), and others in locomotion. In the rostral GRN, excitation of bilateral Chx10 neurons leads to halting of locomotion through inhibition of locomotor circuits (Bouvier et al., 2015), and unilateral excitation leads to turning (Cregg et al., 2020). Recently, two subpopulations of Chx10^ON^ RSNs have been found, with those that project to the cervical cord leading to head orientation and those to the lumbar cord to hindlimb arrest (Usseglio et al., 2020). Given that we found relatively homogeneous properties in Chx10^ON^ RSNs in the GRN, it may be that the different functional subtypes are electrophysiologically indistinguishable.

What is the role of these neurons in behaviour? In terms of movement, the reticular formation – and in particular the GRN – has been studied primarily in terms of “command,” or reticulospinal, neurons. For example, in 1859, Mauthner cells were identified in fish, where they were found to initiate turning movement (Sillar, 2016). Chx10 neurons in the Xenopus reticular formation are involved in initiating bouts of swimming (Li and Soffe, 2019). In cats, neurons in the GRN can initiate postural changes or locomotion (see:Mori, 1987), and they have several roles in mice (vide supra).

Optogenetic experiments have been used to sort out the role of reticular formation neurons. But one problem with this approach is that activating a population as a whole may obscure the role of more precisely defined sub-populations. For example, low doses of light would be expected to excite the lower rheobase neurons (interneurons), which would in turn activate the RSNs. Higher doses of light may excite both, but this could conceal the role of discrete sub-populations. One way to parse function would be to record the activity of these neurons during behaviour. It has recently been found that about one half of Chx10 neurons in the GRN are active during locomotor stop events, but even within these neuronal sub-populations, there seems to be some diversity in function: that is, individual neurons are not active during every stop event (Schwenkgrub et al., 2020). Furthermore, some Chx10 neurons were active at the onset of and during locomotion (Schwenkgrub et al., 2020), consistent with some of them receiving input from the mesencephalic locomotor region (Bretzner and Brownstone, 2013). And some are not active during locomotion, but are during grooming, or when the animal is stationary (Schwenkgrub et al., 2020). Similarly, three functional types of glutamatergic neurons were described in the lamprey, in a similar region of the reticular formation (one that receives inputs from the mesencephalic locomotor region). Each of these neuronal types has a distinct function: initiation, maintenance or stopping swimming (Juvin et al., 2016; Gratsch et al., 2019). That is, it is clear that reticular formation (GRN) neurons have diverse functions in behaviour.

Less attention has been paid to local circuitry in the GRN. Local inhibitory circuits could facilitate bilateral coordination, as in Mauthner cells (Shimazaki et al., 2019), or could conceivably aid in tuning descending commands. But the role of local excitatory interneurons is not obvious. Indeed, these neurons could form a positive feedback loop, leading to unwanted oscillatory behaviour.

Perhaps insight on the role of these interneurons could be gleaned through comparison with spinal circuits. In the spinal cord, several populations of excitatory interneurons are necessary for the production of normal coordinated activity, such as locomotion (see: Kiehn, 2016). If the GRN was simply for integration of descending inputs from various movement centres to produced descending commands, then local inhibitory interneurons may be sufficient to tune the commands. But if the GRN plays a role in the selection and/or *generation* of complex movement, then local circuitry – including excitatory interneurons – may be required. It is interesting that activating Chx10 interneurons led to changes in RSN properties: firing rates in RSNs were higher than could be elicited by current injection, suggesting that dendritic inputs could drive these cells into a nonlinear range, as has been observed in other neurons such as spinal motoneurons (Heckman et al., 2003). That is, interneuronal activity could change the nature of the descending commands – for example, sculpting tonic to phasic activity.

Several studies have modeled the internal organization of the reticular formation and its selection of appropriate RSNs for a motor behaviour (Humphries et al., 2006, 2007). It has been suggested that local nodes of small interneurons (although in their case inhibitory) form synapses with reticulospinal neurons, and are organized through connections with other nearby interneurons within their node. This “small-world” network was hypothesized to ensure rapid cross-network synchronization, consistent node selection for a particular motor action, and increased persistent activity that facilitates ongoing drive required for the appropriate motor response (Humphries et al., 2007). These properties were proposed to be consistent with a role of the GRN in action selection (Humphries et al., 2005),. Although these models used inhibitory interneurons, the concept of small-world networks may be generalizable to the architecture that we have demonstrated here. (The interested reader is directed to the clear analysis of Humphries et al., 2007, for further reading.)

In summary, we have shown that there are at least two distinct subtypes of Chx10^ON^ neurons in the GRN: a local interneuronal and a reticulospinal population. And we have shown connectivity between these, such that the interneurons enhance the firing of reticulospinal neurons. We do not yet know how these circuits function to produce behaviour, although we can propose that these local circuits are important for ensuring that the actions selected by higher motor circuits are processed to ensure that spinal circuits produce those behaviours.

## ACKNOWLEDGEMNTS

We thank Dallas Bennett and Charlotte Ryder-Burbridge for animal care and technical support, and Dr Xiaolu Sun for her preliminary experimental work. This work was funded by the Canadian Institutes of Health Research (RMB: MOP 136981; YZ: MOP-110950) and Wellcome Trust (RMB: 110193). RMB’s position is supported by Brain Research UK. Equipment used for photostimulation experiments was funded by the Canadian Foundation for Innovation (228190, RB) and the Nova Scotia Research and Innovation Trust (RB).

